# Canalized morphogenesis driven by inherited tissue asymmetries in *Hydra* regeneration

**DOI:** 10.1101/2021.12.08.471744

**Authors:** Lital Shani-Zerbib, Liora Garion, Yonit Maroudas-Sacks, Erez Braun, Kinneret Keren

## Abstract

The emergence and stabilization of a body axis is a major step in animal morphogenesis, determining the symmetry of the body plan as well as its polarity. To advance our understanding of the emergence of body-axis polarity we study regenerating Hydra. Axis polarity is strongly memorized in Hydra regeneration even in small tissue segments. What type of processes confer this memory? To gain insight into the emerging polarity, we utilize frustrating initial conditions by studying regenerating tissue strips which fold into hollow spheroids by adhering their distal ends, of opposite original polarities. Despite the convoluted folding process and the tissue rearrangements during regeneration, these tissue strips develop a new organizer in a reproducible location preserving the original polarity and yielding an ordered body plan. These observations suggest that the integration of mechanical and biochemical processes supported by their mutual feedback attracts the tissue dynamics towards a well-defined developmental trajectory biased by weak inherited cues from the parent animal. Hydra thus provide an example of dynamic canalization in which the dynamic rules themselves are inherited, in contrast to the classical picture where a detailed developmental trajectory is pre-determined.

## Introduction

The emergence of a body axis is a central step in animal morphogenesis. In many cases, the animal’s body axis provides two separate guides for establishing the basic layout of the developing body plan; a symmetry axis and a polar direction. The alignment of the axis serves as a skeleton defining a symmetry of the animal’s body plan, while the polarity of the axis marks a breaking of symmetry, guiding the development of functional tissues asymmetrically along the axis. Here we take advantage of the relatively simple body plan of the fresh water animal *Hydra*, to separate these two features of the body axis and focus on the emergence of polarity along a developmental trajectory. *Hydra*’s tubular body is characterized by a single oral-aboral body axis [1]. The alignment of this axis defines the radial symmetry axis of the animal, whereas the polarity of the axis reflects the breaking of symmetry marked by the presence of a head at the oral end of the animal and a foot at the aboral end. The use of live imaging allows us to follow the establishment of the oral-aboral body axis at high spatial and temporal resolutions throughout whole-body regeneration from small excised tissue strips, starting from the initial folding of the tissue towards the emergence of a fully developed mature animal.

The alignment of *Hydra*’s body axis is supported by parallel arrays of supracellular muscle fibers called myonemes that are oriented parallel to the body axis in the ectoderm and in a perpendicular, circumferential orientation in the endoderm [2-4]. This bilayered muscle-like structure provides a skeleton for the animal, supporting its tubular symmetry and allowing its behavioral movements. The polarity of the body axis is manifested at multiple levels, starting from the asymmetric distribution of various signaling molecules along the head-to-foot axis [5-7], and eventually realized in the different morphological features and functional tissues in the animal [1]. In particular, the head organizer localized at the tip of the mouth of the animal has a special role in keeping the integrity of the body plan and continuously defining the polarity of the body axis by constantly inducing position-dependent signals [8]. During whole-body regeneration, after the removal of the existing organizer, a new head organizer has to form to establish and stabilize the emerging body-axis polarity.

A tissue segment excised from a mature *Hydra* has a strong memory of both the alignment and the polarity of the original body axis of the parent animal [4, 9-11]. These inherited features provide important initial conditions that dominate the morphogenesis process and trigger the emergence of a regenerated body axis, stabilizing both its alignment and polarity. An excised tissue segment first folds into a hollow spheroid in order to regenerate [4, 12]. We have previously shown that excised tissues inherit the parallel actin fiber arrays and that this order is partially maintained during the folding process, conferring an initial directionality in the folded spheroids [4]. The alignment of the supracellular actin fibers in excised tissues confers a structural memory that persists during regeneration, and defines the alignment of the body axis of the regenerated animal. We confirmed that this initial directionality and the structural inheritance it conveys are present down to the smallest tissue segments capable of regeneration.

The strong memory of body axis polarity is evident in bisected *Hydra* that regenerate a head or foot according to their original polarity [13]. Later work showed that the memory of polarity is retained even in small excised tissue segments [9-11, 14]. Note that the supracellular actin fibers contain multiple actin filaments of mixed polarity. Thus, while these fibers can support the inherited alignment, they cannot provide a source for polarity memory since these contractile fibers are apolar. The Wnt signaling pathway is known to play a central role in the definition of body axis polarity in *Hydra* [5-7], and local activation of the Wnt pathway can induce head formation [10, 15]. However, despite extensive research, it is still unclear what mediates the memory of polarity in regenerating tissues and specifies the location of the new head organizer. In particular, the initial response following bisection appears similar in head and foot facing wounds [16-18], and the specification of diverse signaling trajectories characterizing head or foot formation is detected only 8-12 hours after bisection [16, 18].

Our recent work provided evidence that the polarity of the emerging body axis can be identified in regenerating tissue segments relatively early after their folding into hollow spheroids by following the emerging defects in the supracellular order of the ectodermal actin fibers [19].

The formation of these defects is an inevitable consequence of a topological constraint that is blind to the axis polarity. Nevertheless, the defect configuration faithfully marks the axis polarity much before the emergence of any morphological features [19]. This result demonstrates that indeed, at least for small tissue segments that fold smoothly into closed spheroids, some information for the axis polarity exists at very early stages of the developmental trajectory, in agreement with the strong polarity memory realized long ago [9], and furthermore that the polarity information is also manifested in structural elements of the supracellular actin fiber skeleton [19].

The polarity of regenerating *Hydra* tissues can also exhibit substantial plasticity and even be reversed under certain conditions, generated e.g. by grafting different tissues or the application of exogenous Wnt [10, 11, 20, 21]. Recent results highlight the importance of the injury response in activating the Wnt signaling pathway associated with oral regeneration [16-18, 22]. However, the injury outcome appears to be modulated by yet unknown signals from the surrounding tissue. In particular, the appearance of a regenerating head at an injury site depends on the tissue context [16-18]. Thus, the developmental trajectory seems to select the location of the emerging organizer among several alternative locations, heavily biased by memory from the parent animal [11].

Advancing our insight into the emergence and stabilization of body axis polarity requires an experimental strategy that can expose regenerating tissue segments to an array of initial and boundary conditions and examine their effects on the developmental outcome. To this end, we adapt a methodology of creating frustrating conditions for the regenerated tissue. We have recently shown that grafting two tissue rings with opposite orientations into a single tissue segment, exposes the plasticity and reorganization capabilities of the regenerating axis polarity with sensitivity also to the original position of the tissue along the body axis of the parent animals [11]. In this work, we further develop this methodology by following the regeneration of rectangular tissue strips. Choosing this initial geometry is motivated by our previous observations that the initial folding of tissue strips creates another type of frustrating initial conditions for the inherited polarity: excised tissue strips fold in a purse-string like manner, bringing together their originally head and foot-facing sides [4]. Any inherited gradients associated with positional information denoting the tissue localization along the body axis of the parent animal would become highly distorted by this folding process (unlike bisected *Hydra* or excised tissue rings where the initial tissue polarity is maintained), yet excised strips regenerate almost exclusively into mature animals with proper morphology [4].

How does a tissue spheroid reorganize following the folding process to regenerate properly along a well-defined body axis? To gain insight into this process, we utilize live microscopy combined with specific markings on the excised tissue strip and follow the tissue deformations and cytoskeletal organization during regeneration. We verified that indeed there is a contact region between the two opposing sides of the tissue strip. Despite this convoluted initial folding step, we find that the regeneration process proceeds along a well-defined trajectory with the tissue deforming in a continuous manner and reorganizing in a way that largely maintains the original tissue polarity. The new head organizer always forms from a region that originated from the head-facing side of the strip, a short distance away from the initial contact site between the two opposing sides of the strip. Similarly, the foot of the regenerated animal forms from a region originating from the foot facing side of the excised strip. The location of the new head organizer coincides with the position of an aster-like defect in the organization of the supracellular actin fibers which emerges early on in the regenerating tissue spheroid. The stereotypical and highly reproducible folding and regeneration process, culminating with the formation of a highly ordered mature *Hydra*, make regenerating tissue strips an excellent model system for further inquiry on the emergence and stabilization of body axis polarity in animal morphogenesis.

## Results

### Folding of an excised tissue strip into a hollow spheroid

Rectangular tissue strips are excised from the gastric (middle) region of mature *Hydra* so that their long axis is aligned with the direction of the body axis in the parent animal (Figs. 1A, S1). The folding dynamics of the tissue strips are followed by time-lapse spinning disk microscopy. We use transgenic animals expressing lifeact-GFP in the ectoderm, to visualize the tissue shape as well as the cytoskeletal reorganization during the folding process. To track specific tissue regions during the folding process and subsequent regeneration, we utilize laser-induced uncaging of a caged dye to stably label particular regions in the excised tissue segment (Methods) [19]. The original polarity of the excised tissue could be trivially maintained if the rectangular strip folded by rolling into a thin cylinder along its long axis, parallel to the parent’s animal body axis and to the supracellular ectodermal actin fibers [9]. However, we find that excised tissue strips do not follow this simple path, but rather undergo a more convoluted folding process that does not fully preserve the tissue’s orientation with respect to the original polarity of the parent animal (Fig. 1A). During the first stage of the folding process, the nearly flat tissue strip curls along its two major axes, parallel and perpendicular to its long axis.

**Figure 1.**
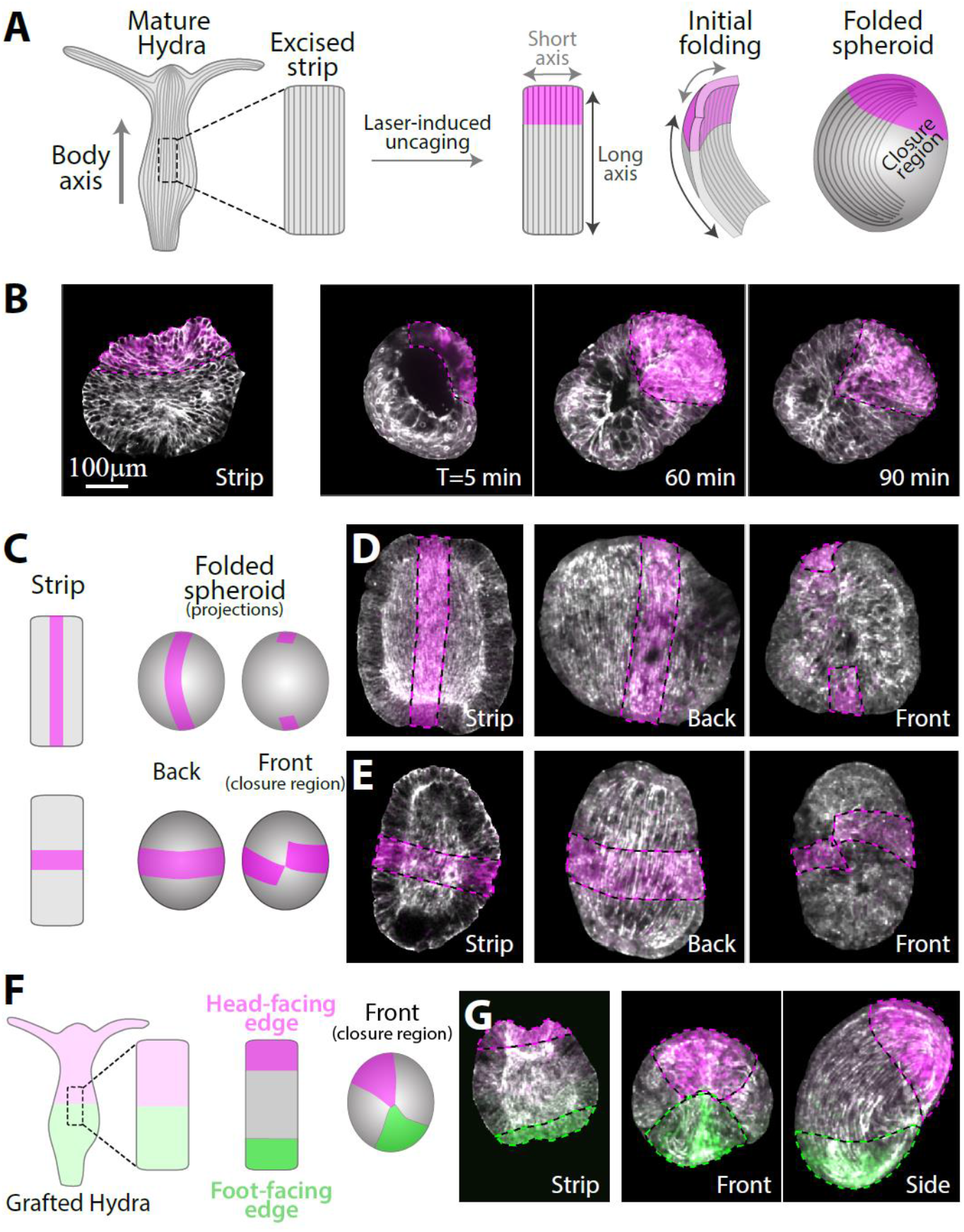
The folding of an excised tissue strip into a hollow spheroid. **(A)** Schematics showing a tissue strip excised from a mature *Hydra* and fluorescently labeled at its head-facing edge. The rectangular strip initially curls along both of its primary axes and subsequently closes into a sealed hollow spheroid. The supracellular ectodermal actin fibers that are aligned with the body axis of the parent animal are shown. (B) Left: Scanning confocal image of an excised strip after laser-induced uncaging of Abberior CAGE 552 at its head-facing edge (see Methods). Right: Spinning disk confocal images from time lapse movies of excised strips labeled at their head-facing edge. The tissue curls along its two major axes and seals with opposite ends of the strip adhering to each other. (C) Schematics and (D-E) images of excised tissue strips that are labeled along a central line parallel to the body axis of the parent animal (C-top, D) or along a medial line perpendicular to the body axis of the parent animal (C-bottom; E). Left: scanning confocal images of the labeling pattern in flattened excised tissue strips. Right: spinning disk confocal projected images of the folded spheroids ∼3 hours after excision from two different viewpoints showing the closure region (front) and the opposite side (back). (F) Schematics and (G) images of an excised tissue strip labeled with two different colors at its edges. The strip is excised from a grafted animal containing Abberior CAGE 552 at its upper half and Abberior CAGE 635 at its lower half, and differentially labeled at both ends by laser induced uncaging. The originally head-facing edge of the strip (magenta) adheres to the originally foot-facing edge (green). All images show an overlay of lifeact-GFP (gray), with uncaged Abberior CAGE 552 (magenta), and in (G) also with uncaged Abberior CAGE 635 (green).

Subsequently, adjacent open edges (brought into proximity by the folding in the first stage) adhere to each other to seal the tissue and form a hollow spheroid (Fig. 1B). In this folding mode, the region that was closest to the head in the parent animal adheres to a region from the opposite end of the strip that was closest to the foot in the parent animal. Naively, this folding pattern seems to generate frustrating initial conditions for the regeneration process since it connects the two tissue edges having opposite polarity with respect to the parent animal. Nevertheless, as shown below, folded strips consistently regenerate into mature *Hydra* with a well-ordered body plan and a normal morphology that largely preserves the original tissue polarity inherited from the parent animal.

To gain a more comprehensive view on the folding process we use different labeling patterns in the excised strip: a central line parallel to the body axis of the parent animal or a medial line perpendicular to the body axis of the parent animal (Fig. 1C). The central line remains nearly parallel to the inherited ectodermal actin fibers in the folded spheroid (Fig 1D, Movie 1).

However, even though the folding process brings together opposite ends of the strip, the edges of the central line do not close on each other (Fig. 1D, right) indicating that the central parts of the originally head and foot facing edges of the strip do not adhere to each other. The medial line remains perpendicular to the actin fibers and nearly closes on itself (Fig. 1E, right). To accommodate these deformations, the tissue stretches in an inhomogeneous manner, with regions near the closure region typically stretching more extensively (and concurrently the bilayer becoming thinner in these regions), whereas regions further from the excised tissue boundaries, in which the supracellular actin fibers remain intact, stretch less (Fig. S2). Repeated experiments with multiple samples (N=24 with central lines, N=8 with medial lines) yielded similar results, indicating that the folding process is stereotypical and reproducible.

To decipher how the originally head and foot-facing edges of the strip adhere to form a closed hollow spheroid that can subsequently regenerate, we label both ends of the strip with two different fluorescent dyes (Fig. 1F). This was done by excising a strip from a grafted animal generated by fusion of two half animals electroporated with different caged probes (Methods) [10]. Laser-induced uncaging at both ends of a strip excised from the middle of the grafted animal differentially labels the strip at the two edges (Fig. 1G). These experiments confirmed that part of the tissue from the originally head-facing edge of the strip always adheres to part of the tissue from the originally foot-facing edge (N=55 samples). Typically, the contact region between the head-facing edge and foot-facing edge is quite small, rather than having an extended interface between the two tissue regions (Fig. 1G). This observation, together with our results utilizing the central and medial line labeling mentioned above (Fig. 1D,E), imply that the tissue strip folds in a somewhat chiral manner bringing into contact two diagonal corners of the initially rectangular tissue. The mechanism driving the tissue segment to complete this complex folding scheme is unknown, presumably involving structural anisotropies inherited in the tissue segment and enhanced by the free open edges. Once folded, the spheroid formed exhibits no apparent initial structural difference in terms of the tissue distribution or cytoskeletal organization between the region that was originally at the head-facing edge or the foot-facing edge of the excised strip.

### Tissue reorganization during regeneration

The excised tissue strips fold into spheroids in a convoluted manner that does not preserve the original layout of the tissue along the body axis in the parent animal (Fig. 1). Nevertheless, excised tissue strips regenerate with extreme fidelity with nearly all samples regenerating into animals that have a normal morphology and a well-defined axis [4]. Given that the original polarity cannot be trivially maintained due to the folding process, we set out to characterize the tissue reorganization in regenerating strips.

Labeling specific regions within excised tissues by laser-induced uncaging of a caged dye allows us to identify and track different regions throughout the regeneration process and determine how their original position within the excised tissue is related to their eventual location in the regenerated animal. We focus on the edges of the excised strip, labeling a region spanning about a quarter of the strip’s length at either its head-facing edge or foot-facing edge, and follow the redistribution of labeled tissue (Figs. 2, S3). Surprisingly, even though the two edges of the strip adhere to each other during the folding process (Fig. 1G), the labelled tissue regions largely maintain their relative position along the animal’s body axis in the regenerated animal as compared to their original position in the parent animal (Fig. 2); The tissue originating from the head-facing edge of the strip is focused at the oral end of the regenerated animal, whereas the tissue originating from the foot-facing edge is localized toward the aboral end (Fig. 2B-D).

**Figure 2.**
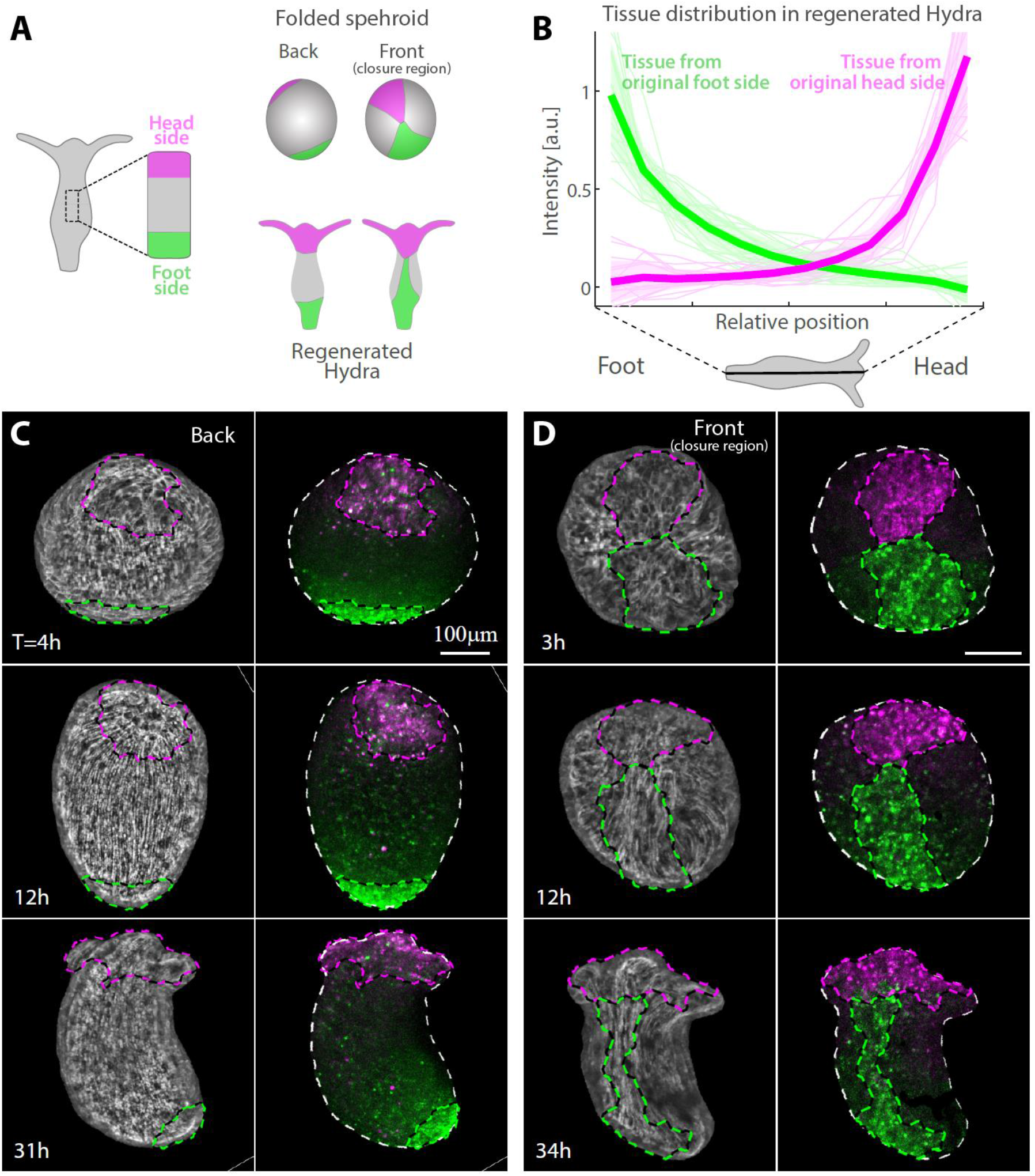
Tissue reorganization during strip regeneration. (A) Schematic illustration of the tissue reorganization during regeneration of an excised strip differentially labeled at its head-facing edge and its foot-facing edge (as in 1F). The regenerating tissue is depicted from opposite viewpoints at an early (top) and late (bottom) stage of the regeneration process. (B) The redistribution of tissue originating from tissue strips labeled either at their head-facing edge (magenta) or foot-facing edge (green) was determined from the relative labeling intensity in the regenerated animals. Graphs showing the relative label intensity along the body axis of animals regenerating from tissue strips labeled at either their head-facing edge (thin lines, magenta; N=19) or their foot-facing side (green; N=19) by uncaging Abberior CAGE 552 (Methods). The position-dependent label intensity is determined from the normalized fluorescence signal intensity within segments along the regenerated animal’s body axis averaged over pairs of projected images of each animal taken from either side (Fig. S3). The population mean of the relative label intensity averaged over all regenerated animals examined (thick lines) and the standard deviation within each population (shaded regions) are also shown. (C, D) Tissue dynamics in regenerating tissue strips differentially labeled at their head-facing edge with uncaged Abberior CAGE 552 (magenta) and their foot-facing edge with uncaged Abberior CAGE 635 (green) in (C) or vice versa in (D) (Methods). Projected spinning-disk confocal images from time lapse movies are shown for a sample imaged from the side containing the closure region (front; D, Movie 3) or the opposite side (back; C, Movie 2). For each time point an overlay of the fluorescent markers labeling the head-facing edge of the strip (magenta) and the foot-facing edge of the strip (green) are shown (Right), together with an image of the supracellular ectodermal actin fibers labeled with lifeact-GFP on the basal surface (gray) overlaid with contours depicting the labeled regions (Left).

The distribution of the originally foot-facing edge is more dispersed, spreading along a larger fraction of the regenerated animal’s body axis and typically exhibiting a tongue of tissue emanating from the foot region toward the head region (Fig. 2D), as compared to the tissue originating from the head-facing edge that is more concentrated at the regenerated head region. Notably, this pattern of tissue deformation is consistent among different samples (Fig. 2, Movies 2-4), indicating the stereotypical nature of the folding and subsequent regeneration.

The dynamics of the tissue reorganization during regeneration are examined by tracking labeled tissue regions over time (Fig. 2C,D; Movies 2,3). Tracking tissue strips differentially labelled at their originally head-facing and foot facing edges using two different probes (as in Fig. 1G), reveals how the seemingly conflicting observations that the head-facing edge and foot facing edge adhere to each other (Fig. 1G) yet the localization of tissue from either edge is largely maintained in the regenerated animal (Fig. 2B), are reconciled. We find that while the tissue in the folded spheroid deforms in a continuous manner, these deformations are not uniform across the tissue, enabling it to mostly retain its original organization along its long axis despite its initial folding along this axis (Figs. 2, S3; Movies 1-3). This arrangement is compatible with the known dominance of the supracellular ectodermal actin fibers in generating the mechanical forces that are essential for the initial folding as well as for the regeneration process [3, 4, 19].

The extent of tissue deformation in the initial closure region is significantly larger than at the opposite side where the actin-fibers organization is stably preserved (Figs. 2C,D; Movies 2,3). Furthermore, the tissue deforms in an anisotropic manner, with the tissue originating from the foot-facing edge of the strip becoming more extended along the new body axis, compared to the tissue originating from the head-facing edge of the strip which remains more localized near the new head (Fig. 2). In particular, the meeting point of the head-and foot-facing edges eventually ends up being located below the head region on one side of the tubular gastric region of the regenerated animal, adjacent to tissue from the originally foot-facing side that is stretched in a highly non-isotropic manner along the new body axis near the meeting point (Fig. 2D, Movie 3). This asymmetry in the tissue spreading between the sides originating from the head and foot-facing ends of the excised strip seems to arise as part of the dynamics of the regeneration process, as the shape of the two labeled edge regions appear similar in the folded spheroid at early stages (Figs. 1G,2C,D).

### Establishment of a new body axis and actin fiber reorganization

Following the characterization of the non-trivial tissue dynamics in regenerating tissue strips, we set out to examine how the new body axis of the regenerated animal is established. Prior to excision, the tissue strip is part of the parent animal which has a clear polar organization along its oral-aboral body axis. We have previously shown that excised tissue pieces retain a parallel array of supracellular actin fibers inherited from the parent animal, that eventually define the alignment of the axis of the regenerating animal [4]. This can be directly visualized by following excised strips in which the tissue is labeled within a line along the oral-aboral axis (Fig. S4B; Movie 1). The labeled line remains aligned with the parallel array of ectodermal actin fibers throughout the regeneration process, and eventually also with the new body axis formed along this direction. Similarly, in excised strips in which the tissue is labeled along a medial line, perpendicular to the original oral-aboral axis, the medial line localizes to the gastric region of the regenerated animal and remains roughly perpendicular to ectodermal fibers that are aligned with the regenerated animal’s body axis (Fig. S4A).

Given that the excised strip does not fold into a cylindrical shape aligned with the original body-axis, but rather folds primarily along its long axis, the alignment of the regenerated body axis with the actin fibers does not define the location of the regenerated head, which in principle could be anywhere along the direction defined by the fibers. The position of the head is determined by the establishment of a new head organizer [8], which is a crucial step in *Hydra* regeneration. We have previously shown that an aster-like defect in the organization of the ectodermal actin fibers marks the location of the new organizer early on in the regeneration process [19]. This aster-like defect remains stationary relative to the underlying tissue, so the defect site reveals the location that becomes the new organizer well before the appearance of any morphological feature there.

Where does the new organizer form in regenerating strips? Time lapse imaging of excised tissue strips expressing lifeact-GFP and labeled at their edges (as in Fig. 1G, 2C,D) allows us to follow in detail the organization of the supracellular actin fibers and identify the defect site marking the location of the new organizer. Initially, as shown by us previously [4, 19], the folded spheroid loses the ordered fibers in the closure region while retaining the ordered parallel array of ectodermal actin fibers elsewhere (Fig. 1D,E). An aster-like defect emerges 8 ± 1 hours after excision (N=10, mean ± standard deviation) (Fig. 3, Movie 4). Once formed, the defect remains stable relative to the underlying cells, marking the tissue region that becomes the site of the new organizer in the regenerated animal (Fig. 3, Movie 4) [19]. These observations, relating between aster-like defects and head formation, are aligned with recent observations in regenerating jellyfish where the establishment of mouth structures was shown to occur at aster-like defect sites, which were also shown to be sites of local activation of the Wnt pathway [23].

**Figure 3.**
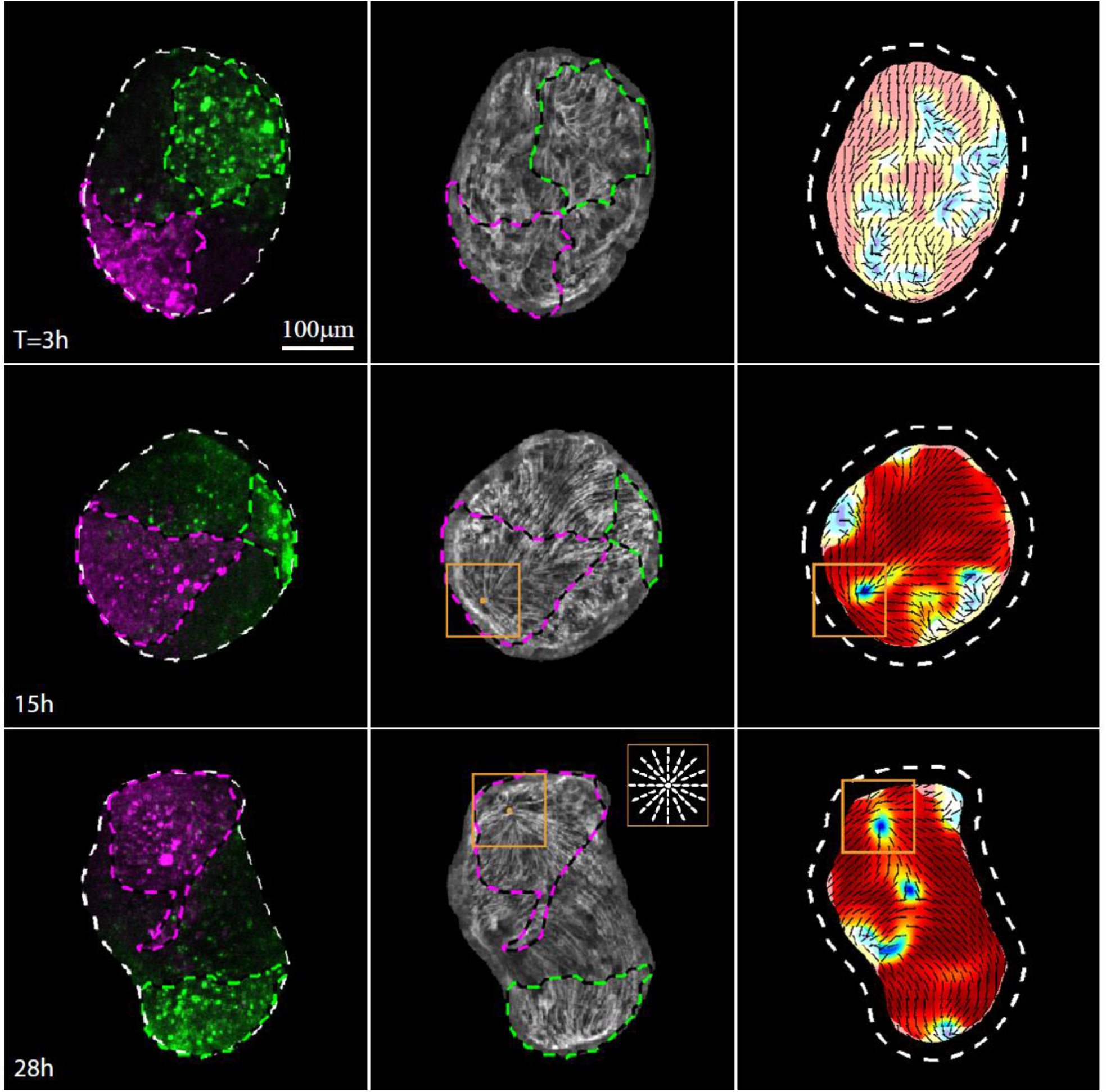
Actin fiber reorganization in a regenerating tissue strip. Images from a time lapse movie (Movie 4) depicting the tissue dynamics and cytoskeletal organization in a regenerating tissue strip expressing lifeact-GFP in the ectoderm (gray) and differentially labeled at its head-facing edge with uncaged Abberior CAGE 552 (magenta) and its foot-facing edge with uncaged Abberior CAGE 635 (green). For each time point, images depicting an overlay of projected spinning-disk confocal images of the fluorescent markers labeling the head-facing edge of the strip (magenta) and the foot-facing edge of the strip (green) are shown (Left), together with an image of the supracellular ectodermal actin fibers on the basal surface (lifeact-GFP, gray) overlaid with contours depicting the labeled regions (Middle), and a map of the corresponding actin fiber orientation (black lines) and local order parameter (color shading, see Methods [19]) with the tissue outline indicated with a dashed line. The location of the aster-like defect in the organization of the supracellular actin fibers that becomes the site of the new organizer at the mouth of the regenerated animal is denoted by an orange box. Inset: schematics showing the actin fiber organization in an aster-like defect.

The defect forms within the initially disordered region of the closure area, as the supracellular actin fibers reform into an ordered parallel array across the entire tissue apart from a small number of localized defects [19]. While the attachment point between the originally head-facing and foot-facing edge is roughly located at the center of the disordered region, we find that the aster-like defect does not form there. Rather, the aster-like defect emerges within the labeled region originating from the head-facing edge of the strip, a few cell diameters away from the original oral boundary of the excised strip (N=10). The clear asymmetry that leads to the formation of the new head organizer within the tissue originating from the head-facing side of the tissue [9], is indicative of a significant persistent initial asymmetric bias in the tissue that is not apparent from the cytoskeletal organization or initial folding of the spheroid early on.

## Discussion

Our observations show that despite the apparent frustration created by the adhesion between the head-facing and foot-facing sides of an excised tissue strip, the folding process and subsequent regeneration follow a well-defined stereotypical developmental trajectory, eventually resulting in an ordered mature body plan. Importantly, the new organizer emerges within the regenerating strip at a similar location relative to its original orientation in the parent animal, in a highly reproducible fashion. Given the complexity of the folding process and the tissue rearrangements during regeneration, as well as the inherent flexibility of *Hydra* tissues that are capable of forming a new organizer at any site within the excised tissue, it seems unlikely that this reproducibility can only reflect pre-patterned biochemical signal gradients instilled in the tissue or signals emerging from injury sites (which in this case span a considerable portion of the tissue). Rather, the highly reproducible regeneration trajectory suggests that the integration of different mechanisms, involving both mechanical and biochemical processes and supported by their mutual feedback, constrain the tissue and attracts the tissue’s dynamics towards a very well defined trajectory.

An important feature to emphasize here is that despite the inherent animal-to-animal variability and further variations in the initial conditions introduced by the rather crude excision step, the strip regeneration process is highly stereotypical, not only in terms of the final outcome but also in the trajectory taken. We show that the initial tissue folding (Fig. 1), the dynamics of the tissue (Fig. 2) and the cytoskeletal reorganization during regeneration (Fig. 3) are all remarkably reproducible, exhibiting essentially the same patterns in all samples. In particular, the location of the new organizer coincides with the site of an aster-like defect in the organization of the supracellular actin fiber that is apparent already ∼8 hours after excision. The canalized trajectory taken by the system is particularly striking given the frustrating initial conditions which do not preserve the original oral-aboral axis of the parent animal, as clearly illustrated by our observations that the most oral and aboral edges of the tissue adhere to each other (Fig. 1G). Nevertheless, the morphogenesis process is highly constrained toward an ordered body plan.

Superficially, the strong canalization of *Hydra* regeneration seems to contradict its flexibility to regenerate a mature animal with a highly ordered body plan from a wide variety of initial conditions and under constraining boundary conditions. However, the form of strong canalization presented by *Hydra* regeneration, appears to be different from the one envisioned by Waddington long ago and that has been serving since then as a guiding principle in developmental systems [24]. This classical picture of canalization envisions a static landscape with imprinted trajectories that guide the dynamics of a developing system in a program-like manner. We suggest that *Hydra* provide an example of *dynamic canalization* which is of a completely different nature from this classical canalization. Instead of imprinted pre-patterned trajectories, the dynamics itself can stabilize its own attractors for proper development; the dynamical rules are inherited rather than the trajectories themselves. Support for the potential for whole-body regeneration under a wide range of configurations and highly variable internal and external conditions, is actually provided by the ability of the tissue to translate weak inherited cues to converge the dynamic trajectory into a stable attractor. In other words, among the multiple different trajectories possible in regeneration, the ones leading to an ordered body plan are selected by weak inherited cues which provide the system enough information to converge to a stable attractor of the dynamics. From the mechanical side, the tissue folding process driven by internal generated forces, the tissue reorganization and the actin-fiber organization leading to an emerging aster-like defect, must all be coordinated with the underlying biochemical signals to support a stable ordered body plan. The convergence of the regeneration process thus suggests that the mechanical processes and the bio-signaling events associated with axis formation are strongly intertwined, and that the robust development arises through constraints embedded in these dynamics and stabilized by mutual feedback mechanisms [25, 26]. Further work is needed to establish this picture of dynamic canalization in *Hydra* and extend this concept to other organisms.

## Acknowledgments

We thank Gidi Ben Yoseph for superb technical assistance. We thank Nitsan Dahan from the LS&E Imaging and Microscopy Unit for help with the laser-induced uncaging. We thank Prof. Bert Hobmayer for generously providing transgenic *Hydra* expressing lifeact-GFP.

This work was supported by a grant from the European Research Council (ERC-2018-COG grant 819174) to K.K., a grant from the Israel Science Foundation (grant No. 228/17) to E.B..

## Methods

### *Hydra* Strains and Culture

All experiments are performed with a transgenic strain of *Hydra* Vulgaris (AEP) expressing lifeact-GFP in the ectoderm [3] (generously provided by Bert Hobmayer, University of Innsbruck). Animals are cultivated in *Hydra* medium (HM; 1 mM NaHCO3, 1 mM CaCl2, 0.1 mM MgCl2, 0.1 mM KCl, and 1 mM Tris-HCl, pH 7.7) at 18° C. The animals are fed three times a week with live *Artemia nauplii* and rinsed after 4 hr. Regeneration experiments are typically initiated ∼24-48 hr after feeding.

### Sample Preparation and Fluorescent Tissue Labeling

#### One color labeling

High resolution marking of specific locations in excised tissue segments is achieved using laser-induced uncaging of a caged dye (Abberior CAGE 552 NHS ester) that is electroporated uniformly into the parent *Hydra* and subsequently activated in the desired regions [19].

Electroporation of the probe into live *Hydra* is performed using a homemade electroporation setup. The electroporation chamber consists of a small Teflon well, with 2 perpendicular Platinum electrodes, spaced 5 mm apart, on both sides of the well. A single mature *Hydra* is placed in the chamber in 10μl of HM supplemented with 16mM of the caged dye. A 75 Volts electric pulse is applied for 35ms. The animal is then washed in HM and allowed to recover overnight prior to tissue excision.

Tissue segments are excised from the middle body section of an electroporated *Hydra* in the following way: the head and foot are removed by two transverse cuts and a cylindrical tube is formed. The cylindrical tube is cut into ∼4 parts by additional longitudinal cuts to generate rectangular tissue segments. An excised tissue segment is then placed and flattened between two coverslips in order to reduce movement during activation. A region of interest is activated by a UV laser in a LSM 710 laser scanning confocal microscope (Zeiss), using a 20× air objective (NA=0.8). The samples are first briefly imaged with a 30 mW 488 nm multiline argon laser at 0.4-0.8% power to visualize the lifeact-GFP signal and identify the region of interest for labeling. Labeling is then done by laser-induced uncaging of the caged dye using a 10 mW 405nm laser at 100%. The uncaging of the Abberior CAGE 552 is verified by imaging with a 10 mW 543 nm laser at 1%. The labeled segments are put in fresh HM for 2-3 hours to allow the tissue pieces to fold into spheroids before placing in a sample holder for time lapse imaging as described below. The uncaged dye remains within the cells of the labeled region and does not get advected or diffuse away within our experimental time window [19]. Over time we observe a reduction in the fluorescence signal due in part to photobleaching.

For statistical analysis of the localization of tissue originating from the head side or foot side of excised tissue strips (Figs. 2B, S3), a large number of tissue strips are labeled either on their original head-facing or foot-facing side in a region spanning roughly ∼1/4 of the tissue. Mature animals electroporated with Abberior CAGE 552 as described above are anesthetized with 2% urethane in HM and the head or the foot of the animal are removed. The remaining tissue is sandwiched between two coverslips and a region along the excision wound, perpendicular to the parent animal’s body axis, is labeled by laser-induced uncaging as described above. To label the entire circumference of the excised animal, uncaging is done from both sides of the tissue by flipping the sample. Strips labeled at either their head or foot-facing edge are excised from the labeled samples as described above. The excised strips are put in fresh HM and allowed to regenerate for 3 days. Subsequently, regenerated animals are briefly treated with 2% urethane, sandwiched between two coverslips separated by a 200µm tape spacer and imaged on a spinning-disk confocal microscope from both sides as detailed below.

#### Two color labeling

Differential labelling of cells originating from regions closer to the head and regions closer to foot of the parent *Hydra* within the same excised tissue segment is realized using two different caged dyes (Abberior CAGE 552 NHS ester or Abberior CAGE 635 NHS ester, 16mM in HM solution) in combination with a grafting technique as follows. Mature animals electroporated with either of the caged dyes are prepared as described above. After recovery from the electroporation (overnight) a fused animal is prepared by grafting the lower half of a bisected animal labeled with one dye to the upper half of a bisected animal labeled with the other dye (the two dyes are used interchangeably for labeling the head or foot regions with no apparent difference). The grafting is done similar to [11] by threading the half animals in tandem (while maintaining their orientation) onto a vertical 75 μm-diameter platinum wire which sticks out of a petri dish that is half-filled with 2% agarose (prepared in HM) and layered with additional HM. The lower animal half is threaded on the vertical wire from the foot side. Subsequently, the upper half of the second animal is gently taken by its tentacles and threaded on the wire on top of the first half (both halves are immersed in HM). Although close contact for a few minutes is sufficient for establishing an initial connection between the two ring tissues, significantly longer time is needed to ensure stable fusion. An additional horizontal wire is placed on top of the two halves to hold them together and the tissue is left to adhere for 6 hours on the vertical wire.

Subsequently, the wires are removed and a grafted *Hydra* consisting of two differently labeled halves is formed. Typically, 12-24 grafted animals are generated in one experiment. The samples are transferred to a 24 well dish in HM solution for an additional 14-16 hours for recovery. The grafted animals are visually inspected under a stereomicroscope and only grafted animals with normal morphology and smoothly connected supracellular actin fibers are subsequently used. Tissue segments are excised from the middle of the grafted *Hydra* as described above, leading to the formation of excised tissue segments containing tissue originating from both of the grafted half animals (and hence labeled with different caged dyes). Photoactivation of the caged dyes is done using a 10 mW 405nm laser at 100% as described above. Typically, activation is done in a region near the head side and a region near the foot side of the excised tissue strips. The two activated regions become differentially labeled following uncaging, allowing us to track the dynamics of the tissues originating from the two different regions within the same regenerating animal. The activation of the Abberior CAGE 552 is verified by imaging with a 10 mW 543 nm laser at 1% and the activation of the Abberior CAGE 635 is verified by imaging with a 5 mW 639 nm laser at 5%.

### Microscopy

Time-lapse imaging of the lifeact-GFP signal and the uncaged Abberior probes is done by spinning-disk confocal microscopy on samples embedded in a soft gel (0.5% low melting point agarose (Sigma) prepared in HM), to reduce tissue movements and rotations during imaging sessions. The general characteristics of the regeneration process in this environment are similar to regeneration in normal aqueous media. The samples are placed in a 25-well agarose sample holder prepared in a 50 mm glass-bottom petri dish (Fluorodish) ∼3 hours after excision. The wells are produced using a homemade Teflon 25-pin comb (1.5 mm diameter pins) with 2% agarose in HM. Each sample is placed in an individual well which is filled with 0.5% low melting point agarose (Sigma) prepared in HM and layered with ∼3-4 mm of HM from above.

Spinning-disk confocal z-stacks are acquired on a spinning-disk confocal microscope (Intelligent Imaging Innovations) running Slidebook software. The lifeact-GFP is excited using a 50mW 488nm laser and the activated Abberior CAGE 552 and Abberior Cage 635 are excited using a 50mW 561nm laser and 100mW 640 nm laser, respectively. Imaging is done using appropriate emission filters at room temperature, and acquired with an EM-CCD (QuantEM; Photometrix). Long time lapse movies of regenerating *Hydra* are taken using a 10× air objective (NA=0.5).

Time lapse imaging is initiated after spheroid folding (∼3 hours after excision) and continues for 3 days at a time interval of 10-15 minutes. Final images of the regenerated *Hydra* are taken after 3 days, after relaxing the animals in 2% urethane in HM for 1 minute and sandwiching them between two glass coverslips with a 200 μm spacer between them. The final images are taken from both sides (by flipping the sample).

### Image analysis

The image analysis tools used for extracting the 2D surface geometry of the tissue, generating 2D projected images of the tissue, and determining the nematic director field are based on custom-written code, as well as adaptation and application of existing algorithms, written in Matlab, Pyton and ImageJ as detailed below.

### Layer analysis

The apical and basal surfaces of the ectoderm are computationally identified in 3D fluorescence z-stacks of the lifeact-GFP signal in regenerating *Hydra* acquired with a z-interval of 3-5 μm using the “Minimum Cost Z surface Projection” plugin in ImageJ (https://imagej.net/Minimum_Cost_Z_surface_Projection). The cost images are generated by processing the original z-stacks using custom-code written in Matlab. First, the signal from the ectoderm layer is manipulated to make it more homogenous within the layer without increasing its thickness, by applying the built-in Matlab anisotropic diffusion filter.

Subsequently, we employ a gradient filter to highlight the apical and basal surfaces as the top and bottom boundaries of the ectoderm layer. The apical and basal surfaces are determined using the minCost algorithm (Parameters used: Rescale xy: 0.5, rescale z: 1, min distance: 15µm, max distance: 45µm, max delta z: 1, two surfaces). The surfaces given by the minCost algorithm are then smoothed by applying an isotropic 3D Gaussian filter of width 1 pixel (after rescaling to an isotropic grid matching the resolution in the z-direction) and selecting the iso-surface with value 0.5.

### Image Projections

2D projected images of the ectodermal actin fibers in the basal surface or the cellular cortices in the apical surface of the ectodermal layer are generated by extracting the relevant fluorescence signal from the 3D spinning disk confocal image stacks based on the smoothed basal and apical surfaces determined above. To obtain the projected 2D image value for each x-y position, we employ a Gaussian weight function in the z-direction with a sigma of 3 μm, which is centered at the z-value determined from the smoothed basal or apical surface with a small (2-3 pixels) fixed offset found to optimally define the desired surface. The signal intensities in the projected images are further normalized employing CLAHE with Matlab’s “adapthisteq” function with Rayleigh distribution and a tile size of 20 pixels.

2D projection of spinning-disk confocal images of the uncaged tissue label are generated by taking a maximum intensity projection of the 3D image stacks in the z-region between the smoothed apical and basal surface determined above.

### Analysis of tissue redistribution in regenerating strips

The redistribution of tissue during regeneration is characterized by following labeled regions in regenerating strips labeled either on their original head-facing or foot-facing side, along regions spanning roughly ∼1/4 of the tissue on either end (Fig. S3C). The localization of labeled tissue in the regenerated animals is deduced from the fluorescence signal intensity distribution. Each animal is imaged from two opposite sides (Fig. S3A). The signal intensity in each side is taken as the 2D mean projection of 3D volumetric imaging. The signal intensity is determined from the 2D mean projection after subtracting the auto-fluorescence signal determined as follows. The contour of the animals (excluding the tentacles) is manually extracted and the auto-fluorescence level in each image is defined from the intensity values within the unlabeled region of the tissue. The labeled region is identified by automated thresholding using Renyi’s entropy method in ImageJ and the auto-fluorescence signal is defined as the 30^th^ percentile of the intensity values outside the labeled region. This measure of the auto-fluorescence is taken since it is less sensitive to the thresholding accuracy. For each image, a centerline along the body axis of the animal is determined using CellTool [27], between manually defined points denoting the tip of the head and the foot of the regenerated animal. The body of the animal is divided into 12 segments, equally-spaced along the centerline (Fig. S3B). The auto-fluorescence corrected mean intensity in each segment is determined and plotted as a function of the relative position along the body axis (defined based on the relative area fraction of the segments). For each animal, the distributions obtained for either side are averaged together (Fig. S3D). The mean intensity profiles are normalized so that the integrated intensity along the relative body position is equal to ¼, reflecting the approximate original area fraction of labeled tissue in the excised strip.

### Actin Fiber orientation analysis

The local orientation of the supracellular ectodermal actin fibers is described by a director field, which is oriented along the mean orientation determined in a small region surrounding every point in space. The nematic director field is determined from the signal intensity gradients in the 2D projected images of the ectodermal actin fibers. as described in [19], and briefly described below.

All the relevant length scales below are given in pixels, with a calibration of 1.28 µm/pixel. The signal intensity gradients 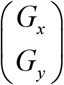 in the normalized images are calculated by convolving the adjusted image, *G*(x,y), with the gradient of a Gaussian with a width of 0.5 pixel. The covariance matrix of the gradient is then calculated from the signal intensity gradient (after averaging using a Gaussian filter with σ=5 pixels), as:

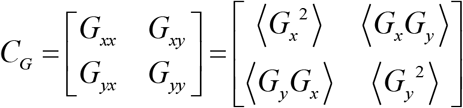

The raw orientation angle is calculated from the covariance matrix as:

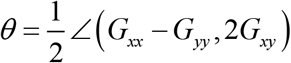

Where <(*x, y*) is defined as:

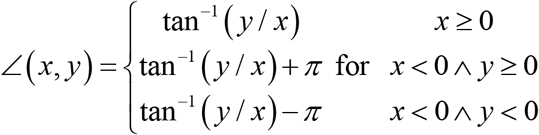

The orientation field is further smoothed by applying a Gaussian filter (σ=3 pixels) on the raw orientation field. The nematic local order parameter, which provides a measure of the local orientational order, is defined as:

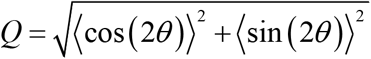

Where *θ* is the smoothed orientation field and the averaging is done over a window size of 32 pixels.

## Supplementary Information

### Supplementary Figures

**Figure S1.**
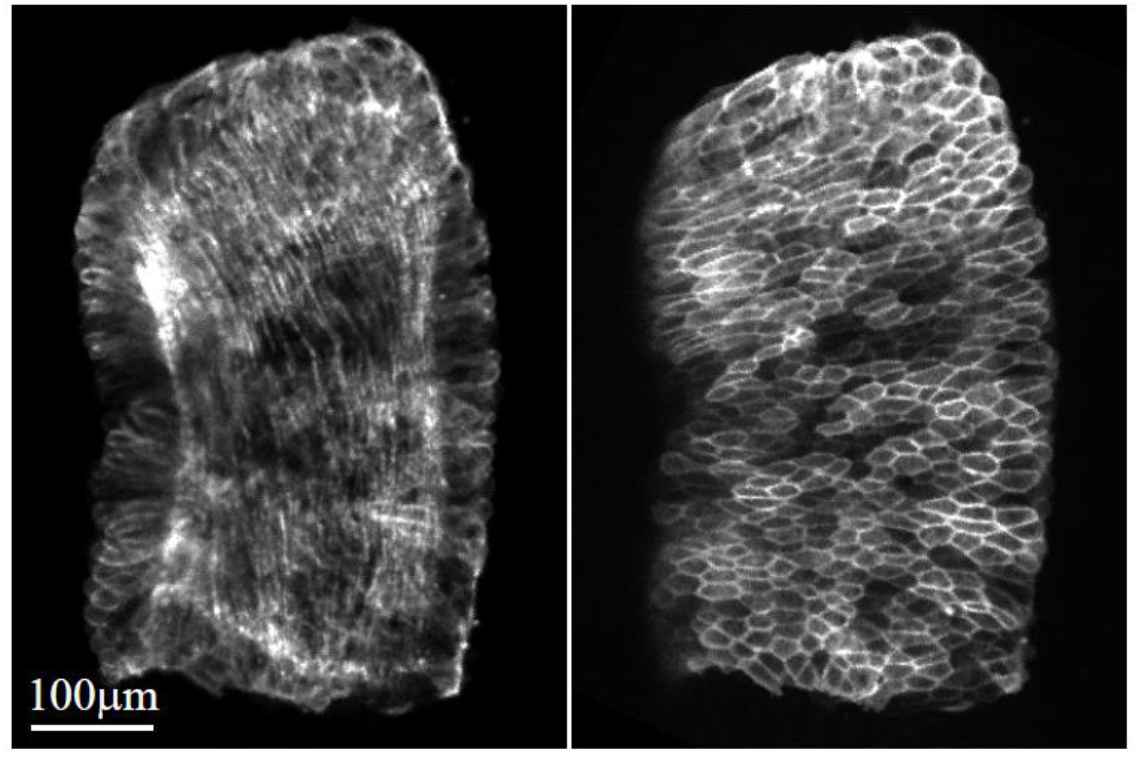
Size and shape of a rectangular tissue strip. Spinning disk confocal projected images of the basal (Left) and apical (Right) surfaces of a rectangular tissue strip excised from the gastric region of a mature *Hydra* expressing lifeact-GFP in the ectoderm. The supracellular actin fibers that are parallel to the body axis of the parent animal are visible in the basal surface of the ectoderm (Left) and the cellular cortices are visible in the apical surface (Right). The length and width of the strips (parallel and perpendicular to the body axis of the parent animal, respectively) is measured by counting cells parallel and perpendicular to the body axis of the parent animal (the size in terms of cell numbers is more reliable since the excised tissue often contracts generating large changes in the apparent size and width of the samples). The measured length and width for a population of excised strips are 36±8 and 16±4 cells (mean±standard deviation, N=11), respectively, for a total of ∼600 ectodermal cells.

**Figure S2.**
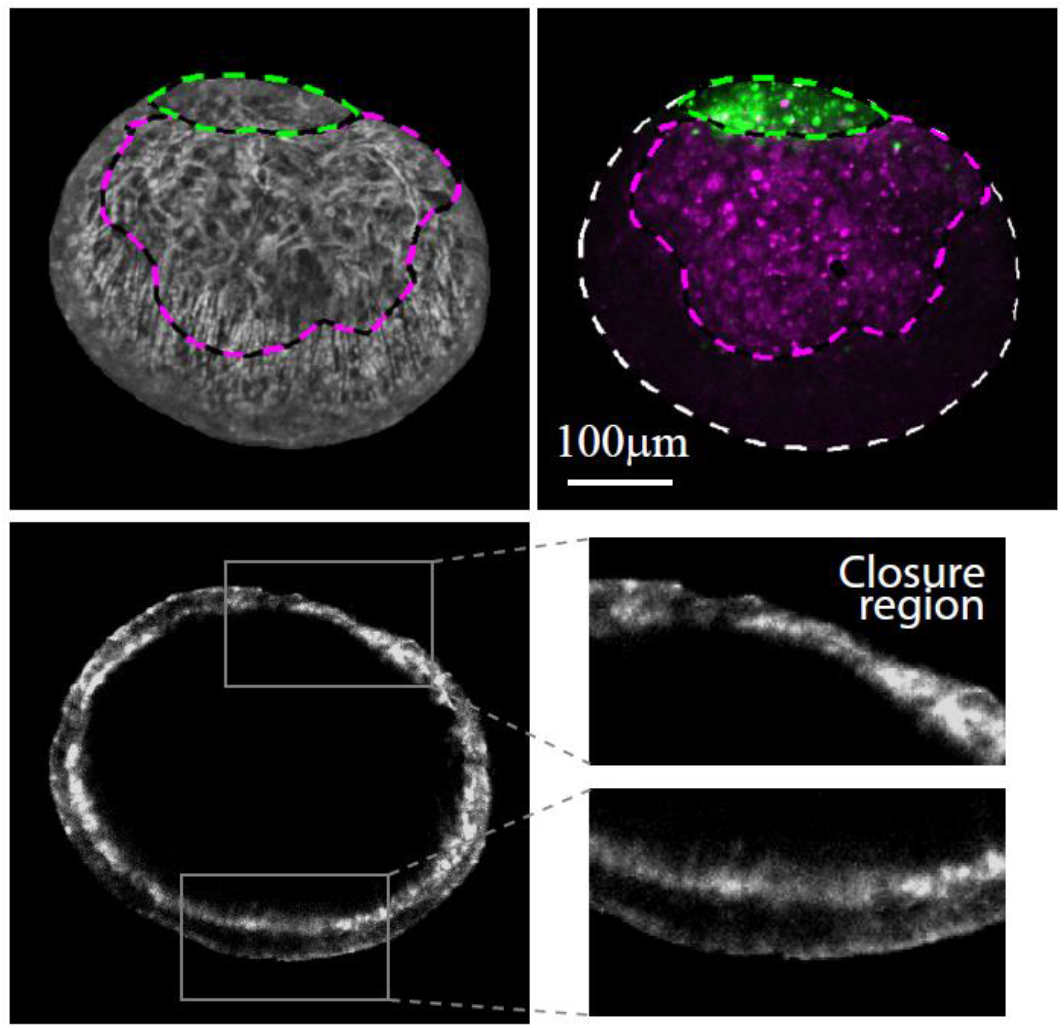
The shape of the initial folded spheroid. Images of a folded spheroid 3 hours after excision, formed from a rectangular tissue strip differentially labeled at its head-facing edge with uncaged Abberior CAGE 552 (magenta) and its foot-facing edge with uncaged Abberior CAGE 635 (green). Top: Projected spinning-disk confocal images of the spheroid imaged from the side, so that the closure region appears in the upper right region of the image. An overlay of the fluorescent markers labeling the head (magenta) and foot-facing (green) edges of the strip are shown (Right), together with an image of the supracellular actin fibers labeled with lifeact-GFP (gray) on the basal surface of the ectoderm overlaid with contours depicting the labeled regions (Left). The actin fibers appear disordered in the closure region (upper half of the spheroid), while the parallel organization of the fibers is maintained away from the closure region (lower half of the spheroid). Bottom: A spinning disk confocal medial cross-section of the tissue spheroid, showing the thickness of the ectodermal layer. The layer becomes thinner in the closure region, due to the tissue stretching that occurs during the folding process, as the tissue seals into a hollow spheroid.

**Figure S3.**
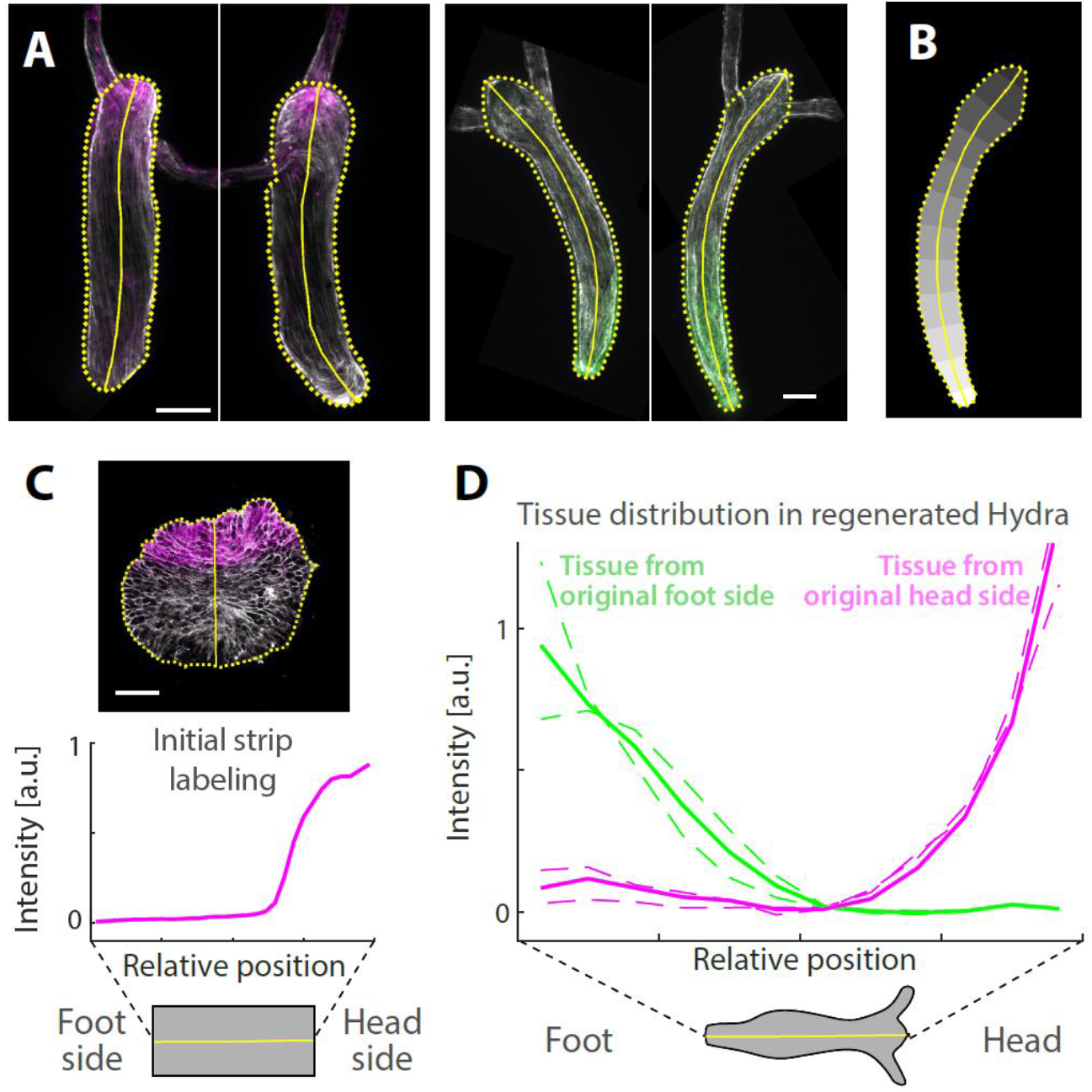
Analysis of the tissue redistribution during regeneration. The distribution of tissue originating from tissue strips labeled either at their head-facing edge (magenta) or foot-facing edge (green) is determined from the relative labeling intensity in regenerated animals. (A) Spinning disk confocal mean projections of the fluorescence signal of uncaged Abberior CAGE 552 for animals regenerating from a strip labeled at its head side (Left, magenta) or its foot side (Right, green). For each regenerating animal, a pair of images are acquired by flipping the sample to visualize the tissue distribution on both sides. (B) A schematic depiction of the definition of the segmented regions used for measuring the position-dependent label intensity along the body axis (Methods). The outline of the animal’s body without the tentacles (dashed yellow line) and the centerline along the body axis (yellow line) are indicated. (C) Top: Scanning confocal images of an excised tissue strip labeled at one end by laser-induced uncaging of Abberior CAGE 552. Bottom: Graph showing the relative label intensity along the excised tissue strip. The images In (A,C) show an overlay of the uncaged label intensity (magenta/green) and the lifeact-GFP signal (gray). (D) Graph showing the relative label intensity along the body axis for the two animals shown in (A), regenerating from a tissue strip labeled on its head side (magenta) or foot side (green), respectively. For each animal, the distributions from either side are shown (dashed lines) together with the average distribution from both sides (thick lines).

**Figure S4.**
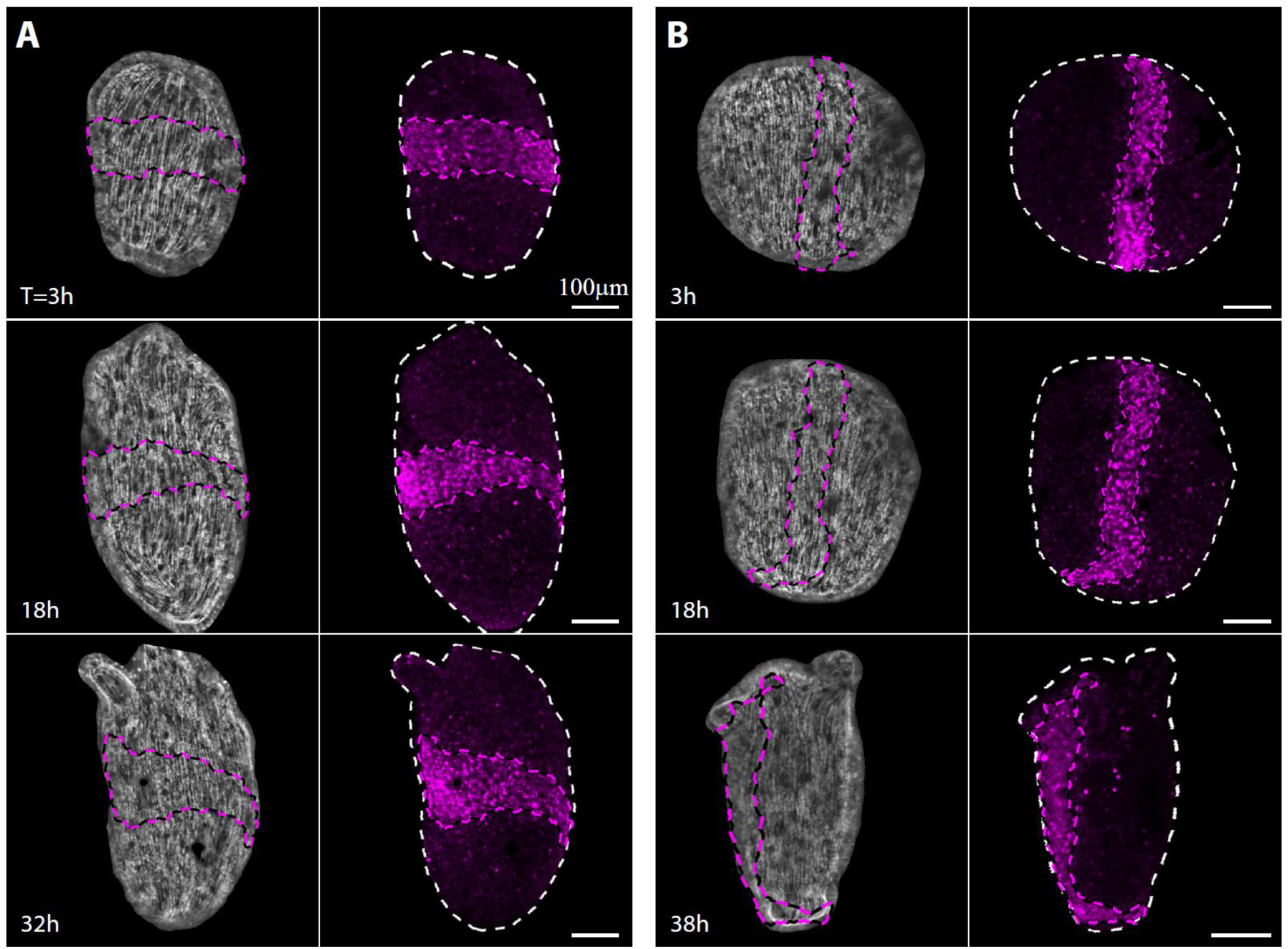
Tissue dynamics and actin fiber organization during regeneration of strips labeled along a medial or central line. Images from time-lapse movies of regenerating tissue strips labeled by laser-induced uncaging immediately following excision along a medial line perpendicular to the body axis of the parent animal (A) or a central line parallel to the body axis of the parent animal (B) (Movie 1). Left: Projected spinning disk confocal images of the supracellular actin fibers in the basal surface of the ectoderm (lifeact-GFP, gray), overlaid with contours of the labeled tissue (magenta). Right: Projected spinning disk confocal images of the fluorescent marker in the labeled tissue originating from the central line in the middle of the strip (uncaged Abberior CAGE 552; magenta). The white dashed line marks the tissue outline. Scale bar is 100 µm.

## Supplementary Movies

**Movie 1. Tissue dynamics in regenerating tissue strips labeled along the central line**. Time-lapse movies depicting the tissue dynamics in regenerating tissue strips expressing lifeact-GFP and labeled along a central line, parallel to the body axis of the parent animal. Upper and lower panels depict two different samples showing the tissue dynamics during regeneration from different viewpoints. The labeled line remains parallel to the ectodermal actin fibers which are aligned with the regenerated animal’s body axis. Left: Projected spinning disk confocal images of the supracellular actin fibers in the basal surface of the ectoderm (lifeact-GFP, gray), overlaid with contours of the labeled tissue (magenta). Right: Projected spinning disk confocal images of the fluorescent marker in the labeled tissue originating from the central line in the middle of the strip (uncaged Abberior CAGE 552; magenta). The white dashed line marks the tissue outline. The elapsed time from excision is displayed (hrs:min), and the scale bar is 100 µm.

**Movie 2. Tissue dynamics in a regenerating strip viewed from its back side**. Time-lapse movie depicting the tissue dynamics in a regenerating tissue strip expressing lifeact-GFP in the ectoderm and differentially labeled at its head and foot-facing edges. The sample is oriented so that the back side of the tissue (opposite from the initial closure region) is visible. Projected spinning disk confocal images of the basal (Left) and apical (Middle) surfaces of the ectodermal layer are shown (lifeact-GFP, gray), overlaid with contours depicting the tissue regions originating from the head (magenta) and foot (green) facing edges of the strip. The supracellular actin fibers are visible in the basal surface of the ectoderm (Left) and the cellular cortices are visible in the apical surface (Middle). Right: Projected spinning disk confocal images of the fluorescent markers labeling the originally head (uncaged Abberior cage 635; magenta) and foot-facing (uncaged Abberior CAGE 552; green) edges of the strip. The outline of the tissue is also denoted (white dashed line). The elapsed time from excision is displayed (hrs:min), and the scale bar is 100 µm.

**Movie 3. Tissue dynamics in a regenerating strip viewed from its front side**. Time-lapse movie depicting the tissue dynamics in a regenerating tissue strip expressing lifeact-GFP in the ectoderm and differentially labeled at its head and foot-facing edges. The sample is oriented so that the initial closure region is visible, showing tissue from the head-facing edge of the strip (magenta) adhered to tissue from the foot-facing edge (green). Projected spinning disk confocal images of the basal (Left) and apical (Middle) surfaces of the ectodermal layer are shown (lifeact-GFP, gray), overlaid with contours depicting the tissue regions originating from the head (magenta) and foot (green) facing edges of the strip. The supracellular actin fibers are visible in the basal surface (Left) and the cellular cortices are visible in the apical surface (Middle). Right: Projected spinning disk confocal images of the fluorescent markers labeling the originally head (uncaged Abberior CAGE 552; magenta) and foot-facing (uncaged Abberior CAGE 635; green) edges of the strip. The outline of the tissue is also denoted (white dashed line). The elapsed time from excision is displayed (hrs:min), and the scale bar is 100 µm.

**Movie 4. Actin fiber reorganization in a regenerating tissue strip**. Time-lapse movie depicting the tissue dynamics and cytoskeletal organization in a regenerating tissue strip expressing lifeact-GFP in the ectoderm and differentially labeled at its head and foot-facing edges. Left: Projected spinning disk confocal images of the fluorescent markers labeling the originally head (uncaged Abberior CAGE 552; magenta) and foot-facing (uncaged Abberior CAGE 635; green) edges of the strip. The outline of the tissue is also denoted (white dashed line). Right: Projected spinning disk confocal images of the supracellular actin fibers in the basal surface of the ectoderm (lifeact-GFP, gray) overlaid with contours depicting the tissue regions originating from the head (magenta) and foot (green) facing edges of the strip. The location of the aster-like defect in the organization of the supracellular actin fibers is denoted by an orange box. The defect forms within the tissue originating from the head-facing side of the strip (magenta), at a site that coincides with the location of the new organizer around the mouth of the regenerated animal. The elapsed time from excision is displayed (hrs:min), and the scale bar is 100 µm.

